# Widespread enhancer co-activity identified by multimodal single cell analysis

**DOI:** 10.1101/2022.10.13.511947

**Authors:** Chaymae Ziyani, Olivier Delaneau, Diogo M. Ribeiro

## Abstract

Non-coding regulatory elements such as enhancers are key in controlling the cell type-specificity and spatio-temporal expression of genes. To drive stable and precise gene transcription that is robust to genetic variation and environmental stress, genes are often targeted by multiple enhancers with redundant action. However, it is unknown whether enhancers targeting the same gene display simultaneous activity or whether some enhancer combinations are more often co-active than others. Here, we take advantage of the recent developments in single cell technology that permit assessing chromatin status (scATAC-seq) and gene expression (scRNA-seq) in the same single cells to link gene expression to the activity of multiple enhancers. Measuring activity patterns across 24,844 human lymphoblastoid single cells, we found that the majority of enhancers associated with the same gene display significant correlation in their chromatin profiles. For 6944 expressed genes associated with enhancers, we identified 89,885 significant enhancer-enhancer associations between nearby enhancers. We found that associated enhancers share similar transcription factor binding profiles and that gene essentiality is linked with higher enhancer co-activity. Our extensive enhancer co-activity maps can be used to pinpoint combinations of enhancers relevant in gene expression regulation and allow us to better predict the effect of genetic variation falling in non-coding regions.

## Introduction

Gene expression regulation is an essential biological process across all organisms and allows for different genes to be activated in a cell type-specific manner, leading to distinct morphologies and cellular functions^1,2^. Gene expression is controlled by genomic regulatory elements such as promoters, insulators, and enhancers^3^. Dysregulation of these elements can lead to a variety of illnesses such as cancer, metabolic syndromes and developmental disorders^4,5^. By harbouring transcription factor binding sites (TFBS), enhancers regulate the spatio-temporal patterns and expression levels of nearby genes irrespective of the position, distance, or orientation relative to the target promoter^6^. To achieve robust expression as well as tight control across cellular contexts, genes utilise multiple enhancers, often with redundant action^7–10^. Indeed, intricate networks of gene expression and regulatory element activity have been revealed in multiple human cell lines^11–13^. In particular, shadow enhancers – sets of enhancers that regulate the same gene, with overlapping activity patterns in space and time – are remarkably abundant and key in controlling developmental gene expression^10,14–16^. Indeed, the action of shadow enhancers has been shown to confer phenotypic robustness to loss-of-function mutations in individual enhancers in loci linked to limb development^10^.

Recent studies have identified enhancers and gene-enhancer links across most human tissues and cell types from ATAC-seq data, ChIP-seq, RNA-seq and CRISPR perturbations^17–19^. However, these studies do not provide information regarding the dynamics of enhancer activity during gene expression regulation and multiple open questions remain, such as whether enhancers targeting the same gene display simultaneous activity, and whether some combinations of enhancers are more often co-active than others. The development of multimodal single cell datasets, particularly those assessing chromatin status (e.g. scATAC-seq) and gene expression (scRNA-seq) in the same single cells^20–22^ now allow us to address these questions at a large-scale.

Here, we exploit the SHARE-seq dataset^20^ with scRNA-seq and scATAC-seq across 24,844 cells in a human lymphoblastoid cells line (LCL) to obtain an in-depth view of enhancer co-activity during gene expression. Starting from cis gene-enhancer associations that we previously identified^23^, for each gene, we correlated the activity levels of all their nearby (within 1 Mb) associated enhancers. Across 6944 expressed genes associated with enhancers, we identified 89,885 enhancer-enhancer associations, amounting to 70.8% of all possible enhancer pairs. Our results confirm the pervasiveness of enhancers with shadow enhancer potential and highlight some of their features such as (i) higher sharing of transcription factor binding sites and (ii) higher enhancer co-activity in essential genes. Our study helps pave the road in our understanding of enhancer co-activity and their role in gene regulation. Knowledge of the relevant regulatory element circuitry, such as which enhancers or combinations of enhancers are relevant for the expression of genes, would allow us to better predict the effect of the hundreds of thousands of GWAS hits falling in non-coding regions.

## Results

### Enhancer-enhancer association predictions from multimodal single cell data

We explore enhancer regulation in a gene-centric way. We have previously exploited the multimodal SHARE-seq single cell dataset^20^, to identify 32,883 gene-enhancer pairs (6944 distinct genes, 7551 distinct enhancers) with correlated activity^23^. Briefly, these gene-enhancer associations were identified using 24,844 LCL cells which contained both scRNA-seq and scATAC-seq data. To focus on enhancer regions, only scATAC-seq peaks overlapping LCL-specific enhancer regions EpiMap repository^19^ were considered. We then correlated gene expression and the activity of nearby enhancers (±1Mb around gene TSS) across cells to identify significant associations (FDR < 5% and absolute Pearson correlation > 0.05, Figure 1a). From 16,463 genes tested, 6944 were associated with at least one enhancer, and 5087 with two or more (max = 35 enhancers, mean = 4.7).

**Figure 1.**
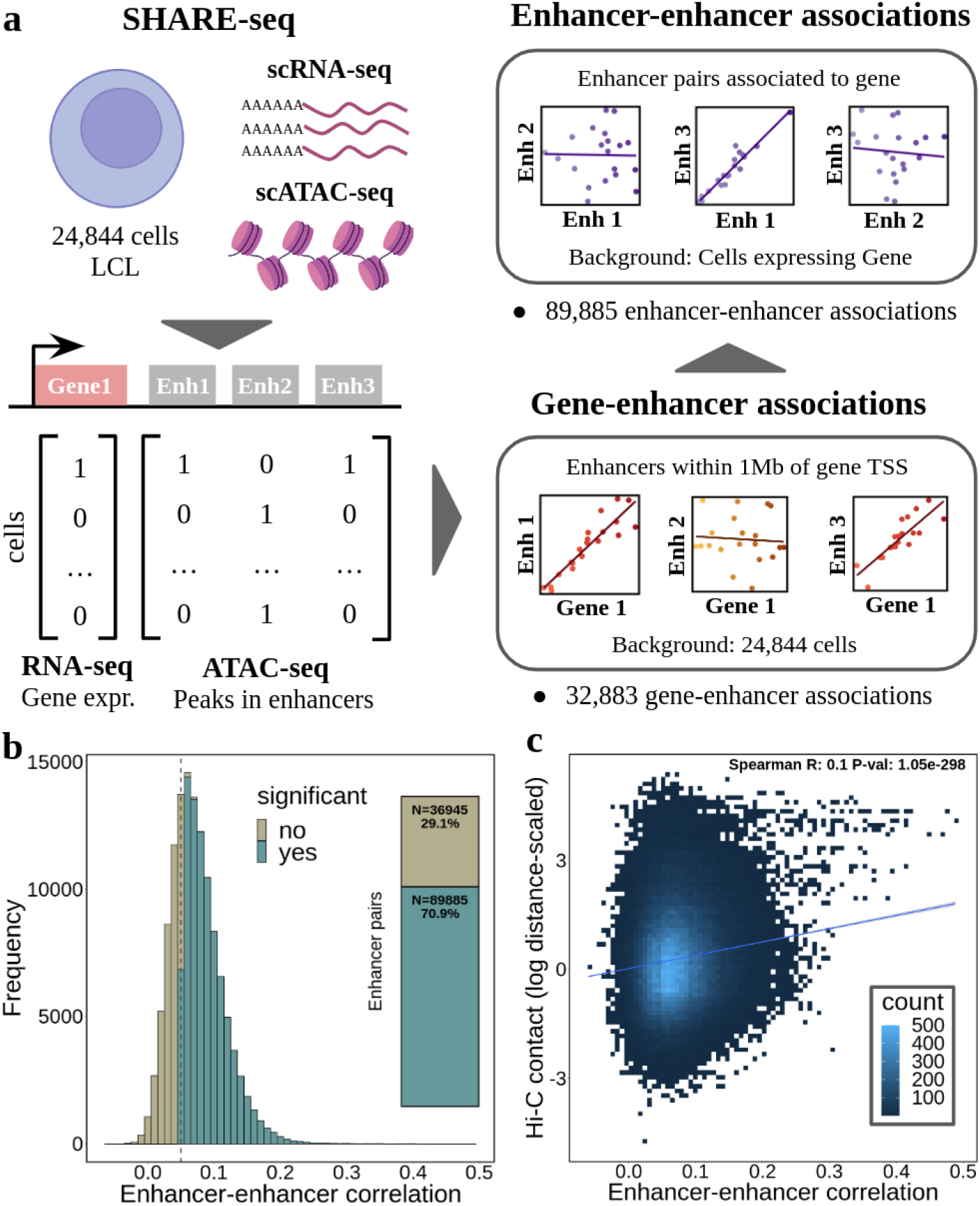
Enhancer-enhancer co-activity overview. **a** scheme of the approach used to determine gene-enhancer associations and enhancer-enhancer associations from SHARE-seq data. Enhancer-enhancer associations were calculated for pairs of enhancers significantly associated with genes. Top left illustration was created with BioRender.com; **b** enhancer-enhancer association correlation distribution (N = 126,830). The inner plot denotes the percentage of significant associations (green colour, FDR< 5% and absolute correlation > 0.05); **c** Hi-C contacts (log distance-scaled, 5kb resolution) per enhancer-enhancer correlation value (N = 126,830).

While several studies identified gene-enhancer associations^17,19,20^, the association between multiple enhancers has not yet been explored at a large-scale. Here, using the same LCL SHARE-seq dataset, we explore the co-activity between enhancers associated with a certain gene. For this, we measure the correlation of enhancer activity (based on scATAC-seq data), for the 5087 genes previously associated with more than one enhancer. Briefly, for each gene we (i) define the set of cells (with RNA-seq and ATAC-seq data) in which the gene is expressed (defined as non-zero expression, mean = 2938 cells per gene), (ii) we gather all enhancer regions associated with the gene (within 1Mb of the gene TSS, mean = 6.1 enhancers per gene), and finally, (iii) for each pair of enhancers, we measure the correlation between their activity (0 or 1) across the cells expressing the gene (see Methods, Figure 1a). Using this approach, we performed 126,830 correlation tests, of which 89,885 (70.9%) were deemed significant (FDR < 5%, absolute correlation > 0.05, Supplementary Data 1). The significant associations comprised 4822 distinct genes and 6743 enhancers and all were positively correlated (Figure 1b). Different significance cutoffs were explored, with 22.1% (FDR 5% and absolute correlation coefficient > 0.1) to 88.6% (FDR 5%, no correlation cutoff) of the tests being deemed significant (Supplementary Fig. 1a). We observed similar proportions of significant enhancer-enhancer associations across cutoffs when considering ABC enhancers and gene-enhancer associations (e.g. 62.9% significant enhancer-enhancer associations with FDR <5% and correlation > 0.05, Supplementary Fig. 1b). We opted for an absolute correlation coefficient cutoff of 0.05 (and FDR 5%) as a moderately strict cutoff, with 70.9% of the tests deemed significant for the SHARE-seq dataset. As a comparison, only 18.4% of the 2,878,013 enhancer-enhancer association tests performed when considering all enhancers within 1Mb of the gene TSS (instead of only the enhancers associated with the same gene) were found significant with the same cutoff (Supplementary Data 2). In fact, only 13.3% of the enhancer pairs were found associated when considering pairs of enhancers not associated with the gene (Supplementary Fig. 2). This indicates that if several enhancers are associated with the same gene, they are more likely to be significantly associated between themselves, as expected.

To confirm the validity of our enhancer-enhancer associations, we analysed publicly available Hi-C data (5kb resolution) for LCLs^24^. We found that the correlation level of the 126,830 enhancer-enhancer association tests correlates with Hi-C contact intensities (Spearman R = 0.1, p-value < 1.1e^-298^, Figure 1c). Moreover, Hi-C contacts between enhancer-enhancer pairs were higher than in distance-matched control regions (Wilcoxon test p-value < 2.2e^-16^, see Methods, Supplementary Fig. 3a). Indeed, 88,283 (69.6%) out of the 126,830 enhancer-enhancer pairs displayed higher Hi-C contacts than expected by their distance (Supplementary Fig. 3b). In addition, when considering a biological replicate with 2788 cells with both scRNA-seq and scATAC-seq (instead of the 24,844 cells used for discovery), we found significant concordance between the enhancer-enhancer correlation levels of the replicates (Spearman R = 0.22, p-value < 2.2e^-16^, Supplementary Fig 4). Together, these results indicate that multimodal single cell ATAC-seq and RNA-seq data can be used to identify enhancer co-activity associations involved in regulating the same gene.

### Prevalence of enhancer co-activity across genes

A key question in enhancer biology is whether enhancers regulate genes in isolation or in simultaneous concert with other enhancers. Enhancer co-activity measurements in single cells can give clues about the cooperativity of enhancers in gene regulation. In our approach, the number of active enhancer combinations observed in a single cell depends on the total number of nearby enhancers associated with a gene. For instance, the gene ABHD4 has three nearby associated enhancers, and we observed all seven combinations of enhancers active in at least a single cell (Figure 2a). The seven combinations comprise (i) three combinations of only one enhancer active in a cell, (ii) three combinations of two enhancers active in the same cell and (iii) one combination with all three enhancers active in the same cell (Figure 2a).

**Figure 2.**
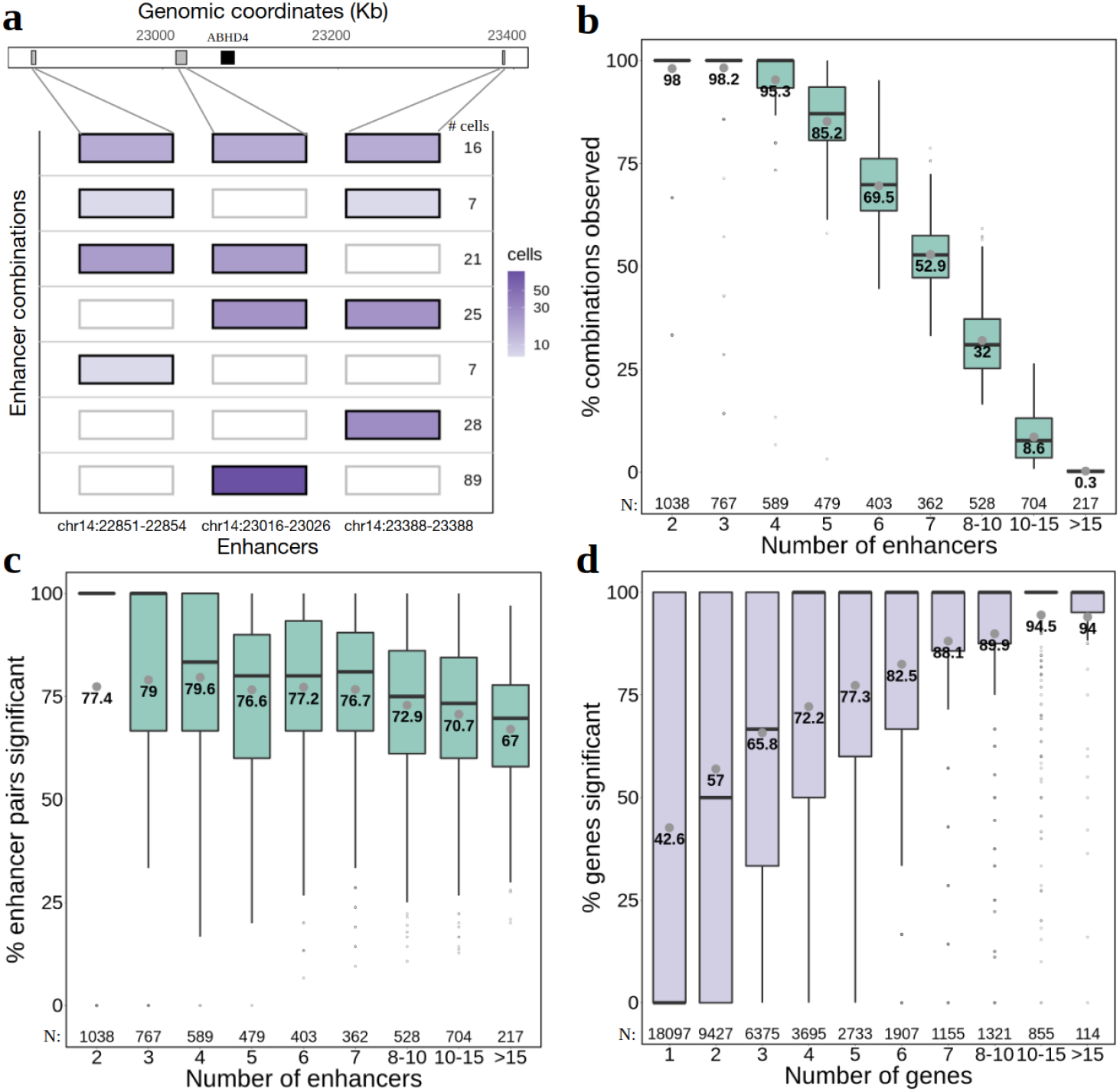
Frequency of co-active enhancers. **a** enhancer combinations observed in single cells for the ABHD4 example gene with three associated enhancers. All seven possible combinations between three enhancers are represented (y-axis), with colour intensity mapped to the number of cells in which the combinations are observed; **b** percentage of enhancer combinations observed in at least one cell (y-axis) per number of enhancers significantly associated with the gene (x-axis). Grey dots and nearby values represent the mean. Sample sizes for each category are provided in the bottom of the plot; **c** Percentage of significantly associated enhancer-enhancer pairs (y-axis) per number of enhancers significantly associated with the gene (x-axis); **d** percentage of genes in which enhancer-enhancer pairs are significantly associated (y-axis) per number of genes in which they were tested (x-axis). A total of 45,679 distinct enhancer-enhancer pairs were analysed. For all bloxplots, the length of the box corresponds to the IQR with the centre line corresponding to the median, the upper and lower whiskers represent the largest or lowest value no further than 1.5 × IQR from the third and first quartile, respectively.

Overall, per gene, we find that on average 76.3% of the possible combinations of enhancers are found to be co-active in at least one cell. Since the number of possible combinations of enhancers increases exponentially with the number of enhancers, we observe a marked decrease in observed combinations with the number of enhancers (Spearman R = −0.89, p-value < 2.2e^-16^, Figure 2b). However, between 95.3% and 98.2% of all enhancers combinations are observed for genes with up to 4 enhancers. While observing combinations of enhancers in at least a single cell provides an idea of the spectrum of possible biological combinations, it does not provide concrete evidence of enhancer co-activity. We then focused on the significant enhancer-enhancer associations (pairs of enhancers instead of combinations). Depending on the number of enhancers per gene, we find that between 67.0% and 79.6% of the possible enhancer-enhancer pairs are significantly associated (Figure 2c, Supplementary Fig. 5). The percentage of significant enhancer pairs decreases with the number of enhancers per gene (Figure 2c, Spearman R = −0.35, p-value = 1.3e^-153^), however, as the increase in number of pairs scales quadratically (and not exponentially as with the combinations) this decrease is less pronounced.

When taking an enhancer-centric perspective, we find that enhancer pairs are more likely to associate with a higher proportion of genes when more genes are present in their vicinity, with as much as 94.5% genes being significantly associated with an enhancer pair when 10 to 15 genes are present in its vicinity (Figure 2d, Spearman R = 0.3, p-value < 2.2e^-16^). This illustrates the presence of genomic regions with high enhancer and gene activity and the high sharing of enhancers across genes, as previously observed^23^. For instance, two enhancers in chr6 (chr6:26104800-26105400 and chr6:26189200-26191000) display significant associations with 21 out of 22 of their neighbouring genes, many of which are found within co-expression gene clusters encoding for Histone proteins^12,25^ (Supplementary Table 1). In summary, we found that enhancer co-activity is highly prevalent across genes, occurring between the majority of enhancers associated with the same gene. This co-activity of enhancers in the same single cells suggests that enhancers do not act in isolation, but rather as a group, possibly functioning as shadow enhancers.

### Molecular features of co-active enhancers

Having sets of co-active enhancers per gene enables the analysis of their properties as a group. We first explored the concordance in transcription factor (TF) binding in co-active enhancers. For this, we overlapped enhancer regions with transcription factor binding sites from ReMap (ChIP-seq data)^26^, obtaining 417,099 enhancer-TF pairs. We then measured the number of distinct TFs with binding sites present in both enhancers of an enhancer-enhancer pair (see Methods). We found that the 89,885 significantly associated enhancer pairs shared higher numbers of TFs (mean = 26.2) than non-significant enhancer pairs (mean = 18.5, Wilcoxon test p-value < 2.2e^-16^, Figure 3a). Moreover, we found that higher enhancer-enhancer correlations correspond to higher number of shared TFs (Spearman R = 0.22, p-value < 2.2e^-16^, Figure 3b). This trend was confirmed when measuring the Jaccard similarity index (JI) between the sets of TFs binding both enhancers in the pair (Spearman R = 0.15, p-value < 2.2e^-16^, Supplementary Fig. 6a). As we found that significantly correlated enhancers were found at moderately lower genomic distances than non-significant enhancer pairs (mean absolute distance significant = 466.6kb, non-significant = 486.5kb, Supplementary Fig. 7), the TF sharing results could be affected by distance. However, we still observe higher TF sharing in 15,242 distance-matched significant (mean = 23.3) and non-significant enhancer pairs (mean = 15.8, Wilcoxon test p-value = 7.7e^-267^, Supplementary Fig. 6b). Importantly, all these results were replicated when using TF data from the MotifMap dataset^27^, which is based on genome sequence scans for known TF motifs (Supplementary Fig. 8). Moreover, similar results were observed when considering ABC model gene-enhancer links and ReMap TF data (e.g. Spearman R = 0.33, p-value < 2.2e^-16^ for the number of shared TFs per enhancer-enhancer correlation, Supplementary Fig. 9).

**Figure 3.**
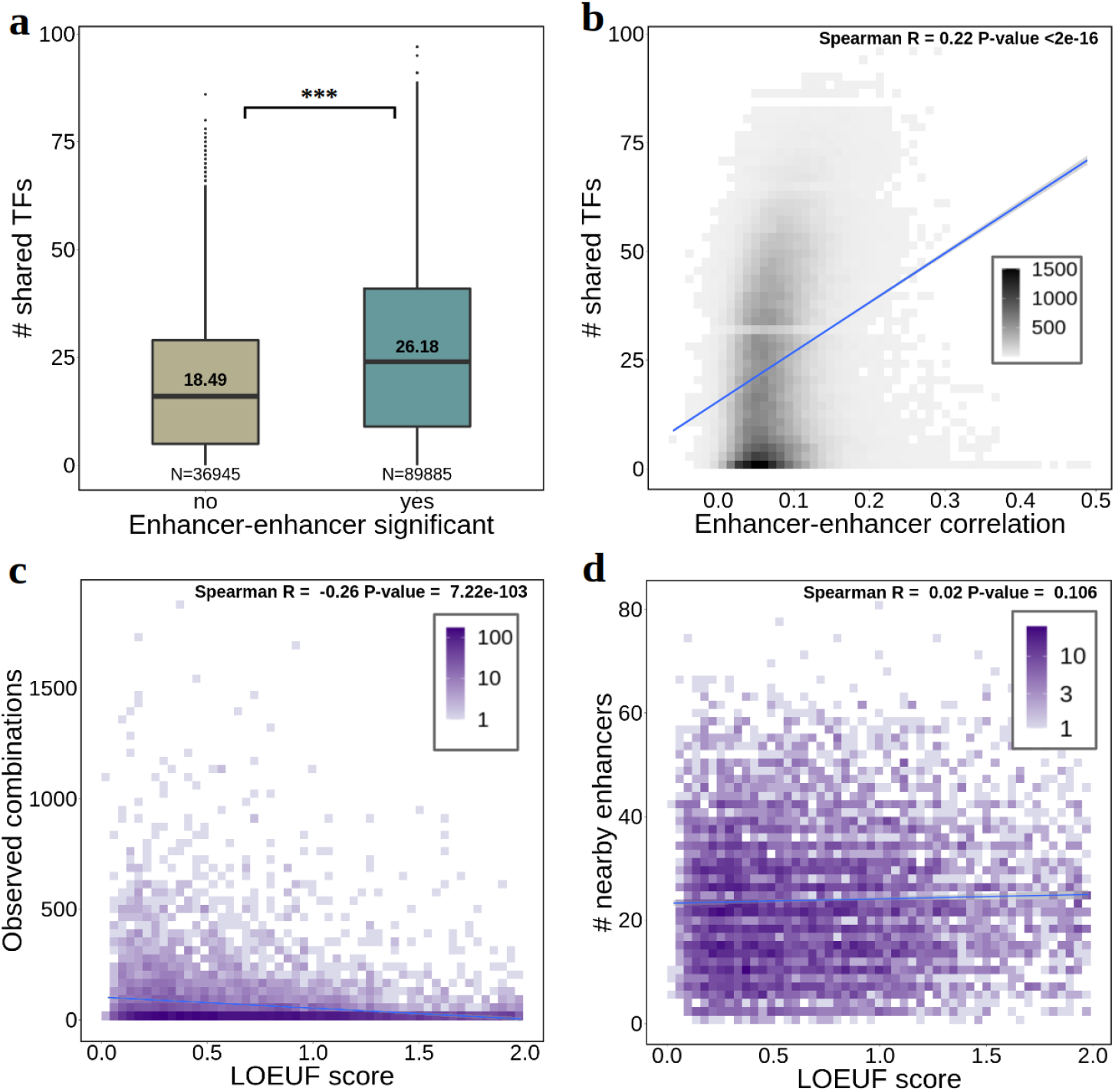
Features of enhancer co-activity. **a** number of distinct TFs with binding sites (ReMap data) in both enhancers of an enhancer-enhancer pair (shared TFs), depending on their association significance. “***” denotes Wilcoxon test p-value < 2.2e^−16^. The length of the box corresponds to the IQR with the centre line corresponding to the median, the upper and lower whiskers represent the largest or lowest value no further than 1.5 × IQR from the third and first quartile, respectively. Values above the median line represent the mean; **b** number of shared TFs per enhancer-enhancer correlation coefficient (N = 126830); **c** number of enhancer combinations observed in at least one cell (y-axis) per gene LOEUF score (x-axis) (N = 6895); **d** number of enhancers within 1Mb of the gene TSS (regardless of gene-enhancer association significance) per gene LOEUF score (N = 6895). Fit lines represent a linear regression model.

Previous studies demonstrated that the number and size of enhancers regulating a gene increases with the gene’s essentiality^7^. We explore this in the context of enhancer-enhancer associations using gnomAD LOEUF scores^28^. For this, we compared the number of significantly associated enhancers and gene essentiality (lower LOEUF scores indicate higher essentiality). We found that gene essentiality is negatively correlated with the number of (i) enhancer-enhancer combinations observed in at least one single cell (Spearman R = −0.26, p-value = 7.2e^-103^, Figure 3c) and (ii) significant enhancer-enhancer pairs (Spearman R = −0.23, p-value = 1.2e^-84^, Supplementary Fig. 10a). This indicates that the more essential a gene is, the more enhancer combinations regulate it. Interestingly, the number of nearby enhancers (regardless of significance) did not correlate with gene essentiality (Spearman R = 0.02, p-value = 0.11, Figure 3d), suggesting that only significantly associated enhancers and their combinations are relevant. Next, we considered enhancer-domain scores from Wang & Goldstein 2020^7^, which reflect the redundancy of a gene’s non-coding regulatory architecture and correlate with gene essentiality. We found significant positive correlation between enhancer-domain scores (higher scores indicate higher redundancy) and the number of (i) enhancer combinations (Spearman R = 0.1, p-value = 1.1e^-15^) and (ii) significant enhancer pairs (Spearman R = 0.08, p-value = 1.0e^-10^, Supplementary Fig. 10b,c). A negative correlation was observed against the total number of nearby enhancers (Spearman R = −0.08, p-value = 5.5e^-11^, Supplementary Fig. 10d). These results highlight the validity of our approach in identifying relevant sets of regulatory enhancers and the importance of robust gene expression regulation through shadow enhancers in essential genes.

## Discussion

While much is known regarding transcription regulation and the potential for multiple (shadow) enhancers to regulate a certain gene^7–9,29^, it is currently unknown whether these multiple enhancers are active at the same time, as this information cannot be obtained from bulk tissue measurements. Our work proposes the use of multimodal single cell RNA-seq and ATAC-seq in the same cells to study the co-activity of enhancers in gene expression regulation. Indeed, by having information of enhancer and gene activity in the same single cell, we were able to define sets of enhancers that are active upon gene expression, and show that enhancer co-activity occurs pervasively across genes. Overall, we found that the set of enhancers that are active upon gene expression can be highly dynamic, with cells presenting disparate patterns of enhancer activity. It is likely that enhancer redundancy serves to drive stable and precise gene transcription, robust to genetic variation and environmental stress^30^. By finding higher numbers of co-active enhancers in essential genes – as well as extensive sharing of transcription factor binding in co-active enhancers – our study corroborates this role of enhancer redundancy. Indeed, we complement previous studies which found a relationship between the number of conserved nucleotides in enhancers and gene essentiality^7^ by showing that this extends to co-active enhancer combinations.

Although single cell data proves useful in connecting gene expression and regulatory element activity^20,31–33^, its usage for determining enhancer co-activity may entail several limitations. First, while we try to expose the breadth of enhancer co-activity that could occur in single cells (e.g. observed enhancer combinations), not finding certain combinations of enhancers in at least one cell does not mean they do not occur, since single cell technology – even when considering >20k cells – may not identify co-active enhancers below certain detection levels. Moreover, some enhancer regulatory patterns may only be revealed under particular cellular contexts or stresses, as has been demonstrated in recent studies of context-dependent quantitative trait loci (QTL) and reporter assays^34–38^. The use of larger datasets of multimodal single cell data, as well as exploring context-dependent effects and other cell-types would likely allow us to observe more enhancer activity combinations. On the other hand, enhancer pair co-activity – even with significant correlation – does not necessarily imply that the two enhancers are active in regulating the same gene or acting together. Indeed, enhancer co-activity could occur as a consequence of the chromatin being open, which in itself could occur stochastically or due to regulation of other nearby genes. Given the high levels of co-expression found between nearby genes^12^ and the sharing of enhancers between co-expressed genes we previously observed^23^, enhancer co-activity is likely influenced by local gene co-expression. Finally, while gene expression and enhancer activity in the same cell are indicative of their relationship, gene transcription is a highly dynamic process and could be partially decoupled in time with nearby enhancer activation^39^, i.e. there could be time lags between enhancer activity and the expression of targeted genes, which would decrease our ability to detect their correlation across single cells.

To exploit known biological knowledge we limited our analysis of enhancer-enhancer co-activity to known enhancer regions from the EpiMap resource^19^ in the same cell line for which single cell data was available (GM12878 LCL). While this allowed us to perform a more focused analysis, we cannot exclude the fact that other genomic regions with enhancer potential were omitted from analysis. In fact, many ATAC-seq peaks in the single cell data fall outside enhancer regions and could be further exploited for regulatory element identification and correlation, as performed in other studies^20,40^. Although we provided an overview of the potential relationship between the several enhancers targeting the same gene, further study is required to understand whether these interact synergistically or even repressively towards gene expression. Moreover, while we found higher bulk Hi-C contacts between co-active enhancers, it is yet unclear if all co-active enhancers interact with each other as well as the gene promoter in a single cell. Further studies with multi-omics single cell datasets, including Hi-C and massively parallel reporter assays, are poised to address these questions in the near future^41,42^.

An improved understanding of enhancer biology would aid the interpretation of non-coding genetic variants. For instance, estimating the robustness of genes to regulatory region mutations can explain why single mutations in their enhancers have little or no effect. Approaches such as regulatory region mutation burden^7,43^ may perform differently depending on the gene regulatory redundancy. In our study, we provide a proof-of-principle framework to define sets of relevant enhancers per gene which can be exploited in gene-trait association testing. Knowing the exact regulatory element circuitry for each gene and accounting for enhancer redundancy is expected to improve the use of whole genome sequencing in the discovery of novel disease genes and in disease diagnosis.

## Methods

### SHARE-seq single cell data

The single cell dataset used in the study was obtained from Ma *et al.* 2020^20^ through GEO (GSE140203). This consisted of preprocessed gene expression counts and ATAC-seq peaks from the single cell SHARE-seq method for the GM12878 lymphoblastoid cell line (LCL). The original dataset included 26,434 genes expressed across 26,589 cells (GSM4156603, rep3, cells with >300 and <7.500 genes expressed) and 507,307 ATAC-seq peaks across 67,418 cells passing quality control (GSM4156592, rep3). On this dataset, we added genomic coordinates (hg19) and Ensembl gene IDs from Gencode v19. We excluded non-protein coding genes, as well as genes in non-autosomes or in the major histocompatibility complex region (MHC, chr6:29500000-33600000). We binarised both the gene expression matrix and ATAC-seq peaks data (values >1 became 1, values = 0 remained 0).

### Gene-enhancer associations

Gene-enhancer association predictions in the GM12878 LCL were obtained from our previous work at Ribeiro *et al.* 2022^23^, Supplementary Data 4. These were identified with the SHARE-seq dataset described above, and also used to identify enhancer-enhancer associations. Briefly, we utilised processed and quality-controlled ATAC-seq peaks from Ma *et al.* 2020^20^ (GSM4156592, rep3). We considered the subset of 24,844 cells that also had gene expression measurements (GSM4156603, rep3). Next, we retained ATAC-seq peaks overlapping GM12878-specific enhancer annotations from the EpiMap repository^19^ (hg19, considering the EnhG1, EnhG2, EnhA1, EnhA2 states from the 18-state chromHMM models), using bedtools (v2.29.2) intersect with the -F0.5 parameter (i.e. requiring >=50% of the peak to be inside the enhancer). Book-ended EpiMap enhancer annotations were previously merged using bedtools merge with default parameters (leading to 33,776 distinct enhancer regions). Finally, we integrated gene expression and open chromatin activity measurements (binarised) for the same cells and enhancer regions within ± 1Mb of a gene TSS were tested for association with the gene through Pearson correlation (equivalent to Spearman correlation when using binary data), in a total of 350,182 tests performed (17,300 distinct enhancers, 16,463 distinct genes). We only considered protein-coding genes in autosomal chromosomes. For each test, we shuffled the expression vector of the gene 1000 times and recalculated the correlation. We then derive an empirical p-value for the probability that the observed value is more extreme than the randomised correlations. To control for the total number of tests we applied the Benjamini–Hochberg procedure for FDR on the empirical p-values. We determined gene-enhancer pairs with correlation coefficient > 0.05 and FDR < 5% as significant gene-enhancer associations (total of 32,883 associations between 7551 distinct enhancers and 6944 distinct genes).

### Enhancer-enhancer associations

We identified enhancer-enhancer associations through a gene-centric approach, using the 24,844 cells from Ma *et al*. 2020 (rep3, GEO:GSM4156592, human LCL GM12878) with both scATAC-seq and scRNA-seq^20^. For this, we started from the 32,883 gene-enhancer associations previously identified with the same 24,844 cells. Then, for each gene, we (i) define the set of cells expressing the gene, which is used as the background of the association test, (ii) for each enhancer associated with the gene, we define the set of cells in which the enhancer is active, (iii) for each pair of enhancers in the set of associated enhancers, we perform Pearson correlation between the enhancer activity vectors across the set of cells expressing the gene. These analyses were performed with custom R (v4.0.4) scripts. We considered 6944 protein-coding genes with enhancer associations to 7551 enhancer regions, performing a total of 126,830 tests. To only consider robust correlation patterns, we excluded 34 genes which were expressed in less than 100 cells. We determined 89,885 enhancer-enhancer associations as significant by having a (i) Benjamini–Hochberg procedure FDR below 5% and (ii) an absolute Pearson correlation coefficient above 0.05, although other cutoffs were explored (Supplementary Fig. 1).

To compare enhancer-enhancer correlation levels between (i) enhancers significantly associated with genes and (ii) enhancers not associated with genes, we performed the same experiment described above, but considering all enhancers in the vicinity of genes (at most ±1Mb away from the gene TSS), instead of only enhancers significantly associated with the gene. For this, 2,878,013 correlation tests were performed and the same significance cutoffs were applied to determine significant enhancer-enhancer associations. For result replication, enhancer-enhancer association tests were also performed for a biological replicate experiment (rep2, GEO:GSM4156591) containing 2788 cells with both scRNA-seq and scATAC-seq data. This provided us data to perform enhancer-enhancer correlation tests in rep2 for 79,788 out of the 126,830 gene-enhancer-enhancer combinations tested previously (in rep3).

To compare our results with gene-enhancer definitions for other studies, we gathered 62,255 gene-enhancer association predictions from Nasser et al. 2021^17^ (ABC model, file: AllPredictions.AvgHiC.ABC0.015.minus150.ForABCPaperV3.txt) for the ‘GM12878-Roadmap’ cell type. Using this dataset, we measured ABC enhancer activity by overlapping SHARE-seq scATAC-seq data as previously. We could evaluate activity for 46,773 associations (10,862 distinct genes, 23,306 enhancer regions) out of 62,256 ABC gene-enhancer associations. From this data, we calculated enhancer-enhancer correlation as previously described and compared multiple correlation and multiple test correction cutoffs to determine significance (Supplementary Fig. 1).

### Hi-C support of enhancer-enhancer associations

We obtained bulk Hi-C data for the GM12878 LCL cell line at 5 kb resolution from Rao *et al*. 2014^24^. We measured KR normalised (MAPQG0) contact between bins encompassing the midpoint of enhancer regions through custom Python scripts. Normalised Hi-C contacts were log2-transformed. Missing data (enhancer-enhancer bins without Hi-C data) was replaced with 0. We then correlated Hi-C contacts with enhancer-enhancer activity correlation levels. To exclude the effect of distance in measuring Hi-C contacts, when indicated in the figure legend, we residualised the Hi-C contact levels for the distance between enhancer-enhancer pairs with a linear regression model. As a control, for each enhancer-enhancer pair, we produced another control pair composed of the first enhancer and an ‘enhancer’ region on the opposite up- or down-stream location in respect to the first enhancer midpoint (e.g. if the first enhancer is at position 5000 and the second at position 8000, the matching control region has the first enhancer at position 5000 but the second at position 2000).

### Transcription factor binding analysis

We analysed two datasets of transcription factor binding sites (i) 4,052,293 binding sites (155 distinct TFs) based on ChIP-seq data collected from ReMap 2022^26^ for the human LCL GM12878 (hg19 assembly) and (ii) 4,474,877 binding sites (445 distinct TFs) based on motif mapping on hg19 from MotifMap^27^. For this last dataset, multiple motifs targeted by the same TFs were combined. To determine sharing of TF binding between pairs of enhancers, we first overlapped TF binding sites with enhancer regions using bedtools *intersect* with -F1 parameter (i.e. ensuring that 100% of the binding site is contained with the enhancer region coordinates). In this manner, we obtained 417,099 TF-enhancer combinations for the ReMap dataset and 46,112 combinations for the MotifMap dataset. For each dataset, we then counted the number of distinct TF with binding sites in both enhancers for each enhancer-enhancer pair analysed (TF sharing). Enhancer pairs without any TF binding site in the dataset were counted as having 0 shared TFs. To exclude a potential bias from the distance between enhancers in their likelihood of sharing TFs, we compared TF sharing between a set of enhancer-enhancer pairs matched for distance. For this, we matched 15,242 non-significant enhancer pairs to 15,242 subsampled significant enhancer pairs with a maximum absolute distance difference of 5% in between the enhancers. For instance, to match a non-significant enhancer pair apart for 1000 bp, we randomly sampled a significant enhancer pair with a distance between 950 and 1050 (1000 ± 1000 * 0.05).

### Gene essentiality analysis

To associate gene essentiality and number of enhancers associated with the gene, we obtained “loss-of-function observed/expected upper bound fraction” (LOEUF) scores per gene from gnomAD v2.1.1^28^ (https://gnomad.broadinstitute.org/). Low LOEUF scores indicate strong selection against predicted loss-of-function variation in a gene, i.e. higher predicted gene essentiality. We correlated LOEUF scores per gene with the number of observed or significant enhancer associations in 6895 genes with an attributed LOEUF score and at least one significantly associated enhancer. We performed the same analysis for enhancer-domain scores (EDS) obtained from Wang and Goldstein 2020^7^, which were available for 6925 genes.

## Data Availability

The enhancer-enhancer associations produced are available for download as Supplementary Data. Code to produce enhancer-enhancer associations and all figures in the manuscript is provided in https://github.com/diogomribeiro/enhEnh. Input data used in this study is available in the public domain. LCL single cell RNA-seq and ATAC-seq (SHARE-seq) processed data is publicly available through GEO (accession: GSE140203).

## Acknowledgements

D.M.R. has been funded by the European Union’s Horizon 2020 research and innovation programme under the Marie Sklodowska-Curie grant agreement No 885998. O.D. and D.M.R. have been funded by a Swiss National Science Foundation (SNSF) project grant (PP00P3_176977). The funders had no role in study design, data collection and analysis, decision to publish, or preparation of the manuscript.

## Author Contributions

C.Z. and D.M.R. performed the experiments and analysed the data. C.Z. and D.M.R wrote the manuscript with inputs from O.D. D.M.R and O.D. designed the experiments, supervised the study and revised the manuscript.

## Supplementary Information

**Supplementary Fig. 1.**
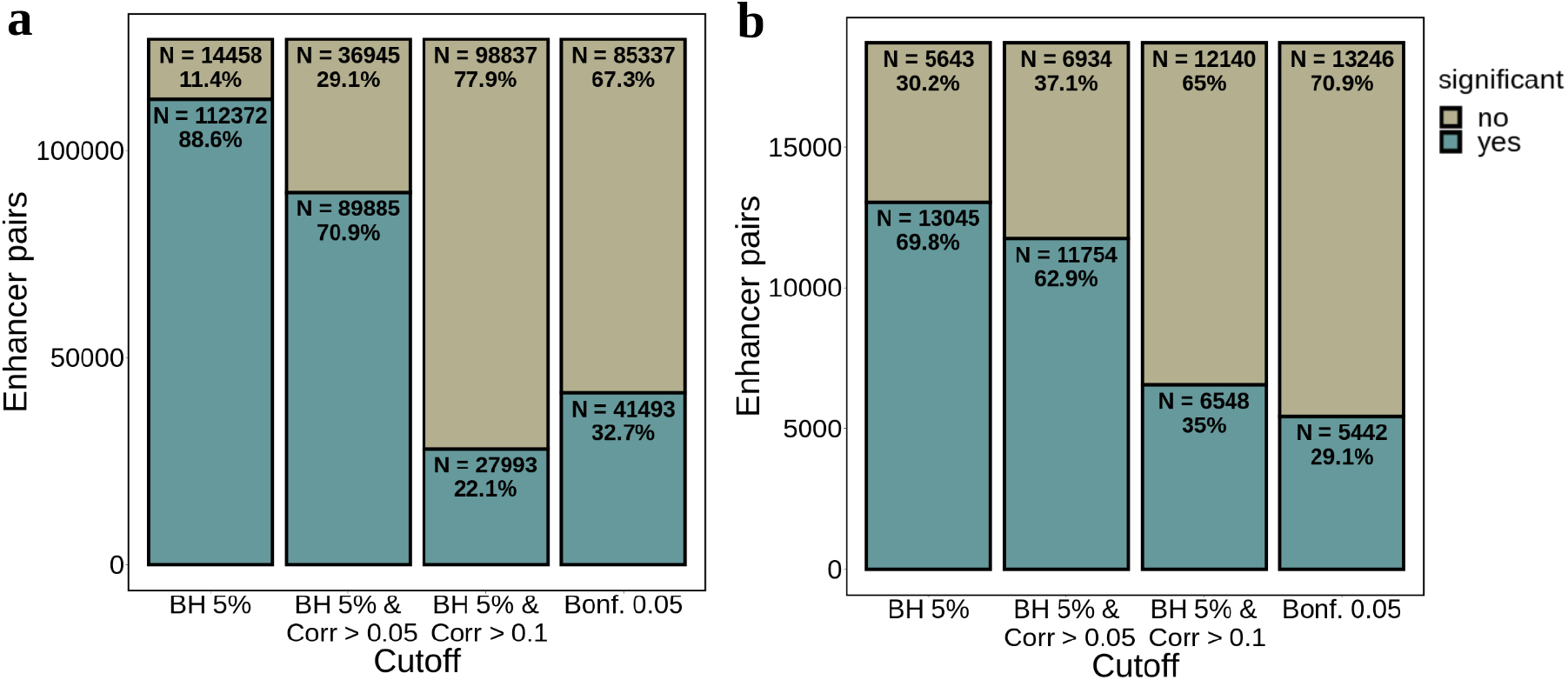
Percentage of significantly enhancer-enhancer associations depending on the multiple testing correction and correlation coefficient cutoff used. **a** enhancer models from EpiMap and gene-enhancer associations based on SHARE-seq single cell data (N = 126,830); **b** enhancer models and gene-enhancer associations based on the ABC model (N = 20,611).

**Supplementary Fig. 2.**
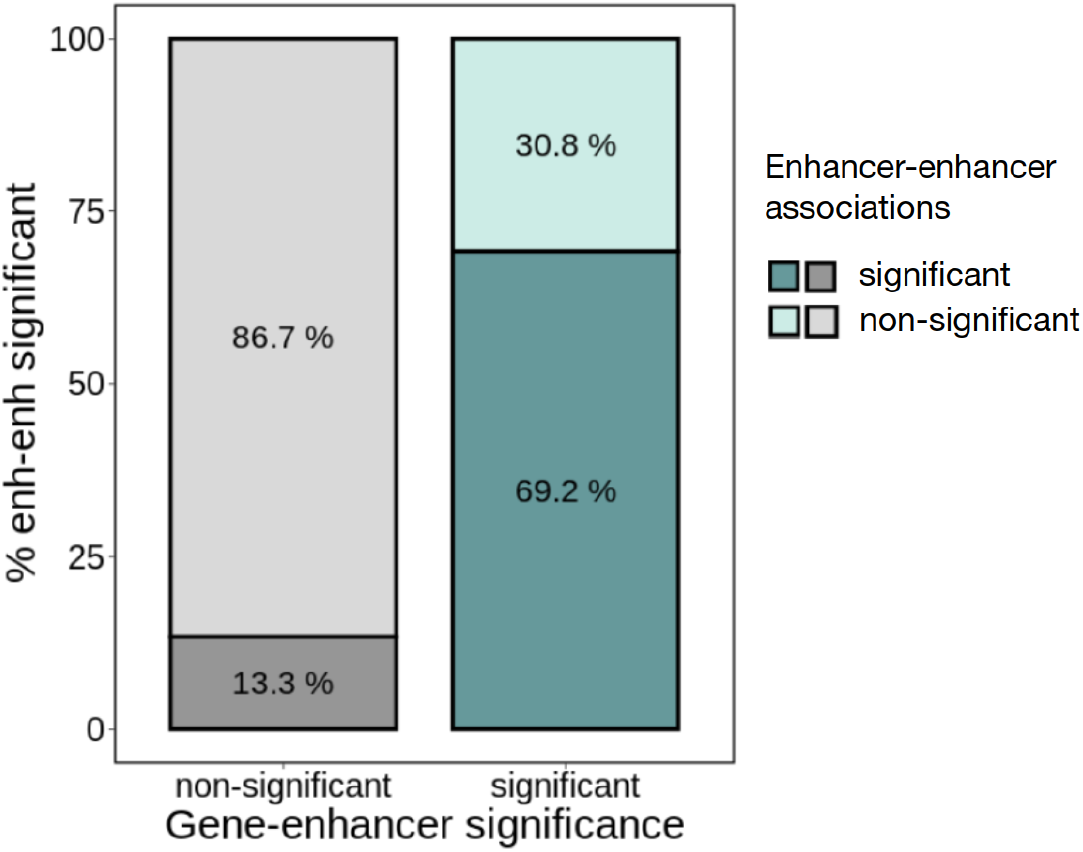
Percentage of significantly enhancer-enhancer associations depending on the significance of the underlying gene-enhancer associations. The gene-enhancer *significant* category refers to cases in which both enhancers are significantly associated with the gene. Conversely, the gene-enhancer *non-significant* category refers to cases where neither of the enhancers is significantly associated with the gene. Results for cases where one enhancer is significantly associated with the gene but the other enhancer is not are not shown. Note that a slightly lower percentage of significant enhancer-enhancer associations (70.9% to 69.2% with significant gene-enhancer associations) is due to the higher burden of multiple test correction in this analysis (N = 2,878,013).

**Supplementary Fig. 3.**
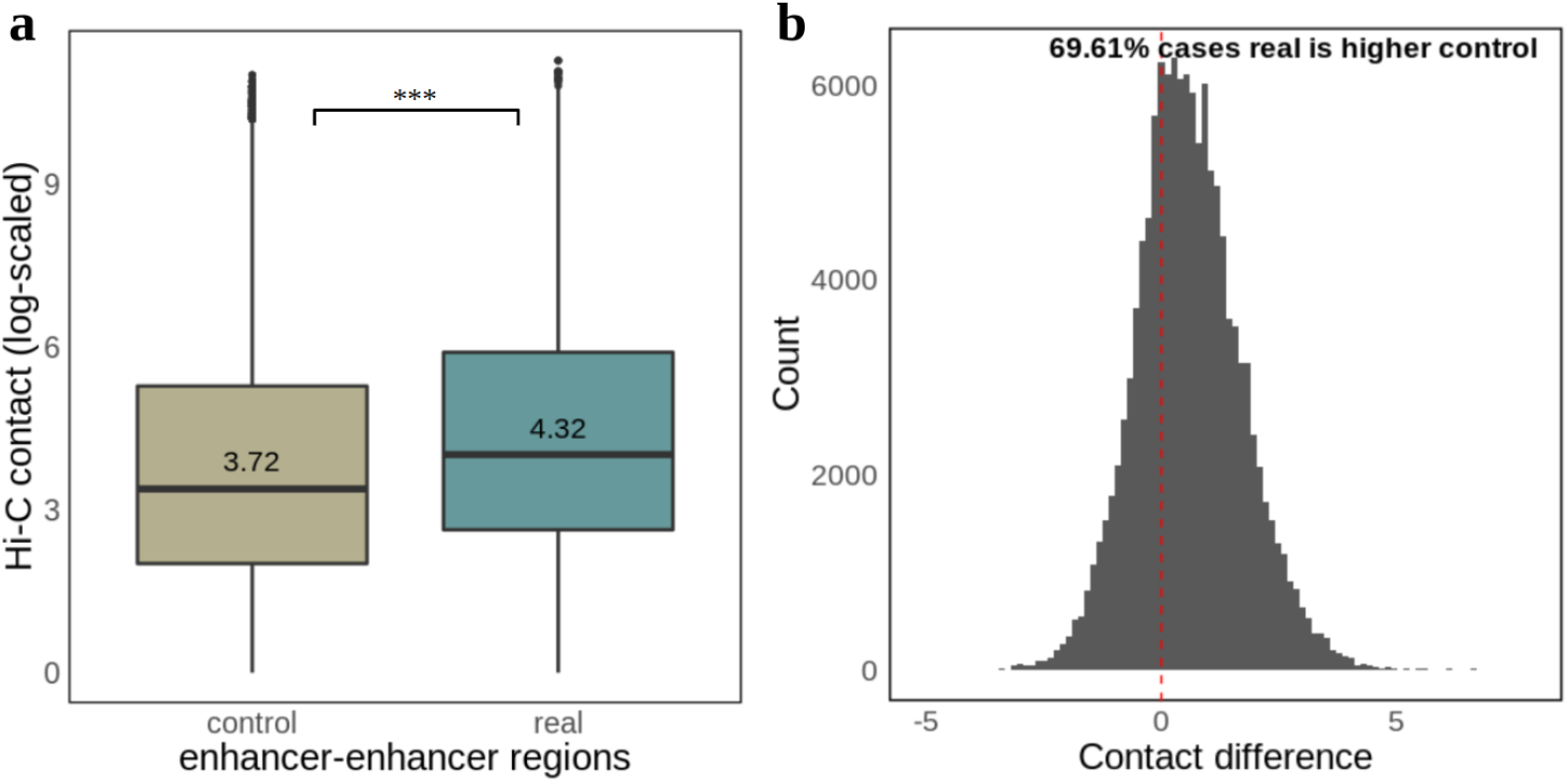
Comparison of Hi-C contacts between real enhancer-enhancer coordinates and distance-matched control regions. **a** boxplot of Hi-contacts (log-scaled) for real enhancer-enhancer coordinates and control regions, in which the coordinates for one enhancer (enhancer 1) are kept, but for the other enhancer are replaced with the upstream/downstream position (see Methods). The length of the box corresponds to the IQR with the centre line corresponding to the median, the upper and lower whiskers represent the largest or lowest value no further than 1.5 × IQR from the third and first quartile, respectively. Values above the median line represent the mean. “***” denotes Wilcoxon test p-value < 2.2e^-16^; **b** difference in contact intensities between real and control regions. A shift of the distribution to the right (above 0) represents higher Hi-C contacts in the real regions compared to control. The midpoint position of the enhancer start and end coordinates was used to calculate Hi-C contacts (N = 126830). Hi-C contact missing data (3.1% of the cases) was replaced with 0.

**Supplementary Fig. 4.**
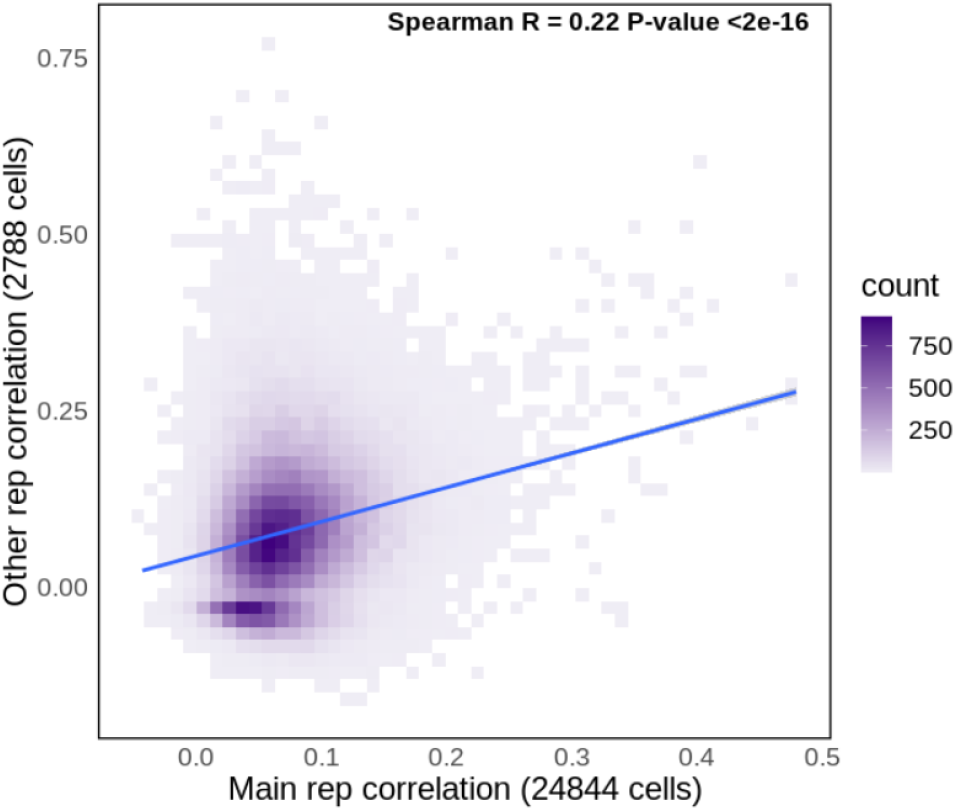
Enhancer-enhancer association correlation values between two biological replicates. Gene-enhancer-enhancer associations tested in rep3 (main rep, 24844 cells, x-axis) were tested for correlation using data from rep2 (other rep, y-axis, 2788 cells). A total of 79,788 gene-enhancer-enhancer associations had sufficient data to be compared (e.g. gene expression in at least 100 cells and ATAC-seq data overlapping the enhancers).

**Supplementary Fig. 5.**
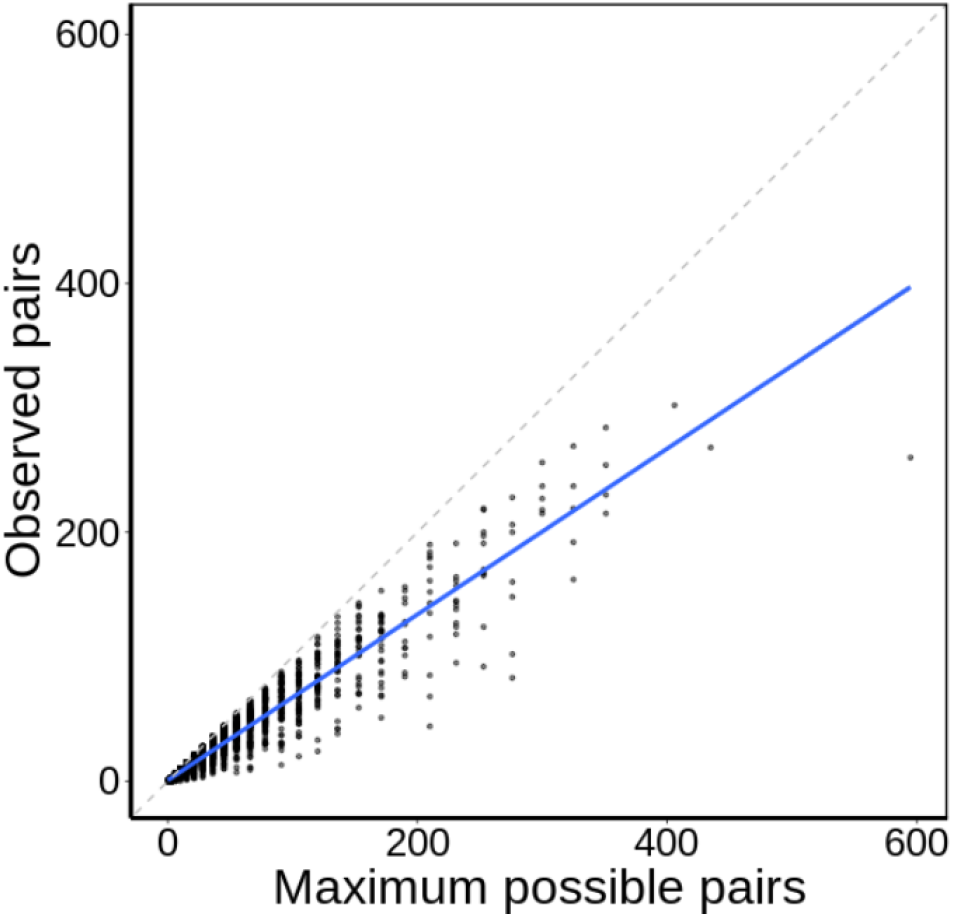
Comparison between observed and maximum possible enhancer-enhancer pairs across genes. Each dot represents a gene. The number of maximum possible pairs scales with the number of enhancers (significantly associated) per gene. The number of observed pairs refers to significant association between the enhancer pair. Fit line corresponds to a linear regression model.

**Supplementary Fig. 6.**
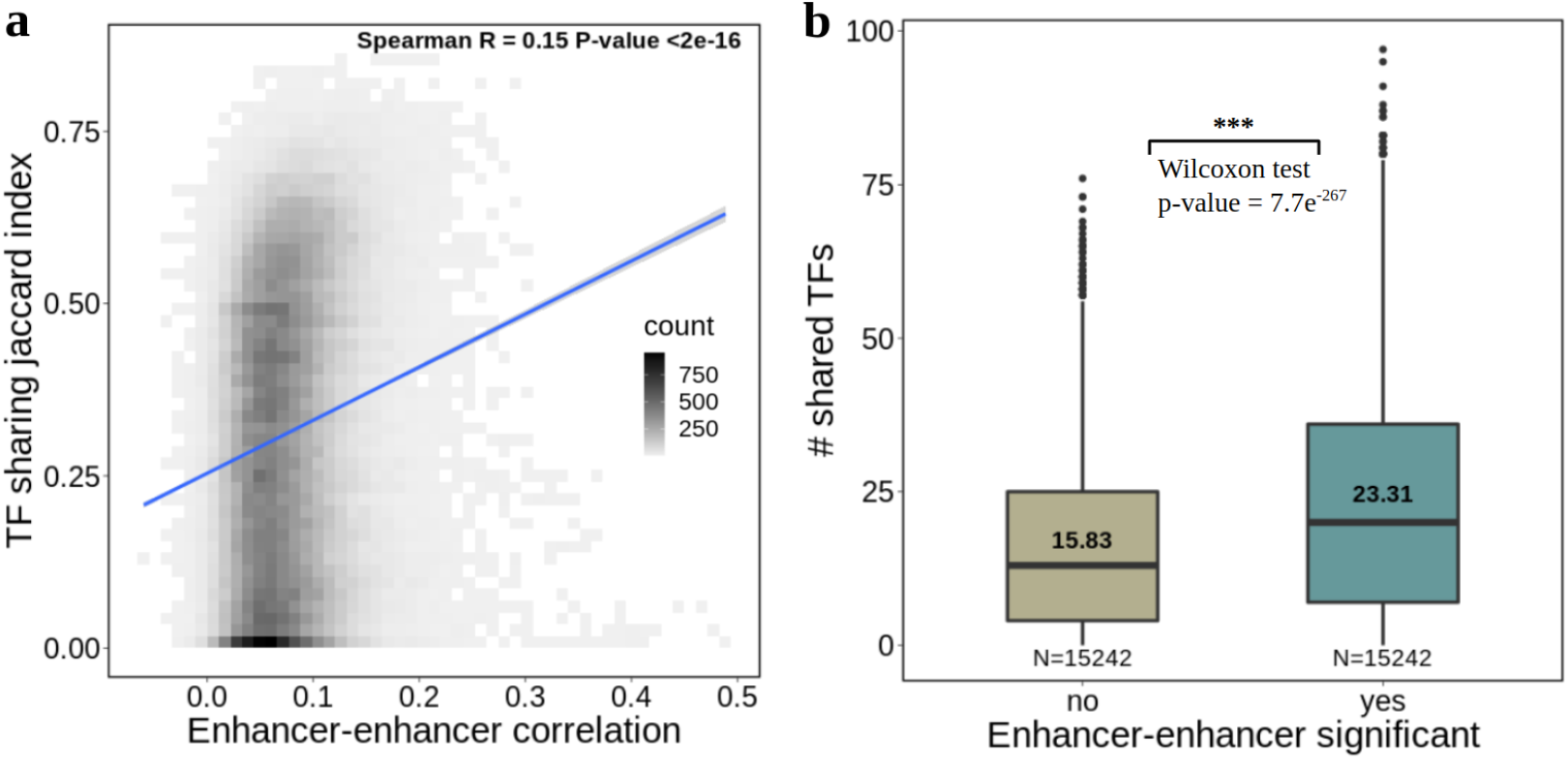
Jaccard similarity index and distance-matched TF sharing using ReMap data. **a** TF sharing Jaccard Index (intersect of TFs found between the enhancer pair, divided by the union of TFs) per enhancer-enhancer correlation coefficient (N = 126,830). Fit line corresponds to a linear regression model with 95% confidence intervals; **b** number of distinct TFs shared between enhancer pairs depending on their association significance. Significant and non-significant enhancer pairs are a subset of all enhancer pairs that are matched for distance (see Methods). The length of the box corresponds to the IQR with the centre line corresponding to the median, the upper and lower whiskers represent the largest or lowest value no further than 1.5 × IQR from the third and first quartile, respectively. Values above the median line represent the mean.

**Supplementary Fig. 7.**
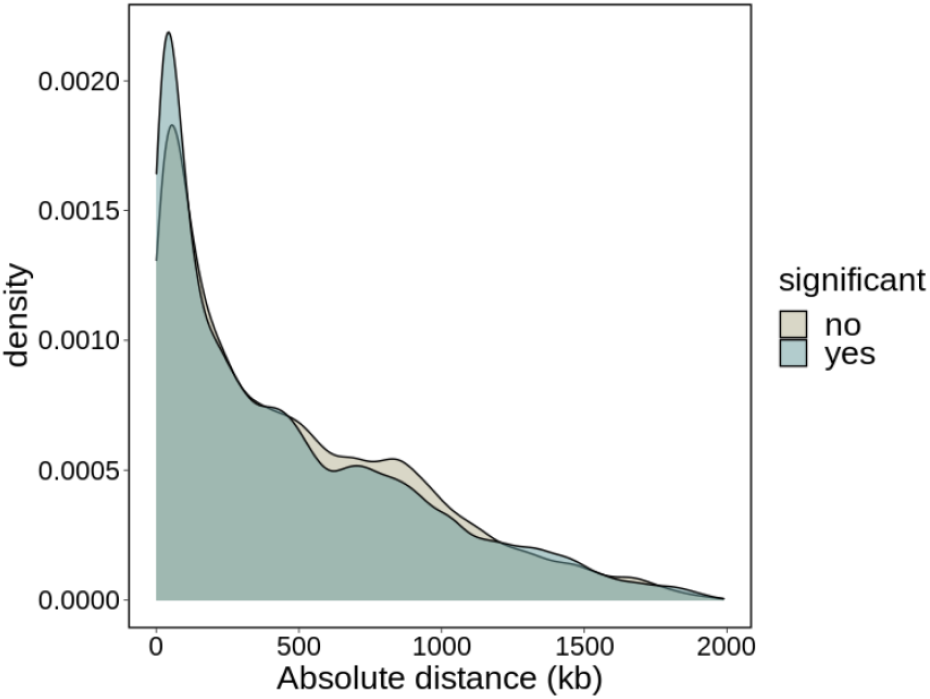
Absolute distance distribution of significant and non-significant enhancer-enhancer associations. The midpoint position of the enhancer start and end coordinates was used to calculate the distance. N = 89,885 for significant enhancer-enhancer pairs and N = 36,945 for non-significant enhancer-enhancer pairs.

**Supplementary Fig. 8.**
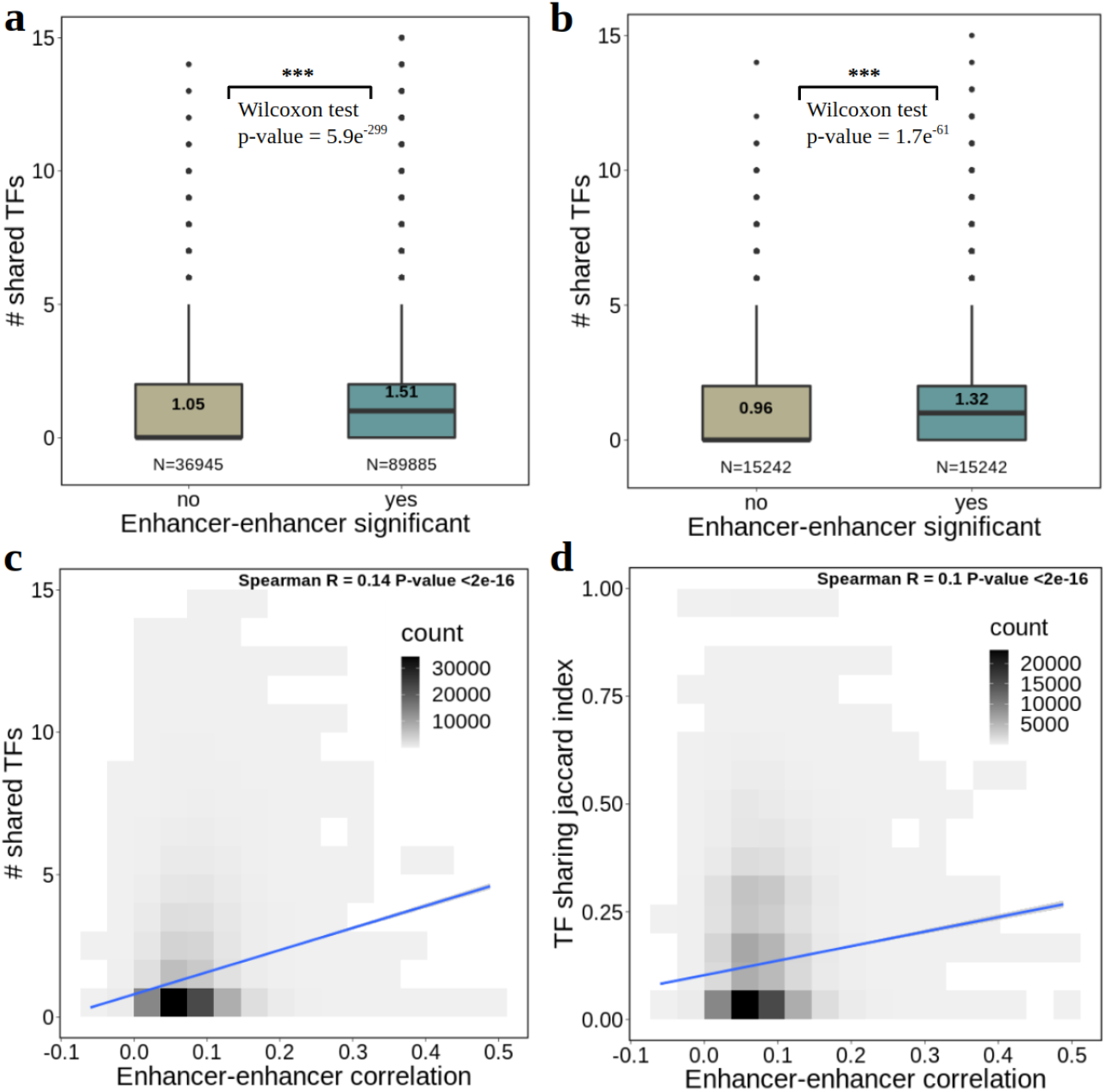
Transcription factor sharing using MotifMap data. **a** number of distinct TFs with binding sites in both enhancers of an enhancer-enhancer pair (shared TFs), depending on their association significance. The length of the box corresponds to the IQR with the centre line corresponding to the median, the upper and lower whiskers represent the largest or lowest value no further than 1.5 × IQR from the third and first quartile, respectively. Values above the median line represent the mean; **b** same as previous, but significant and non-significant enhancer pairs are matched for distance (see Methods); **c** number of shared TFs per enhancer-enhancer correlation coefficient (N = 18,688). Fit line corresponds to a linear regression model with 95% confidence intervals; **d** TF sharing Jaccard Index (intersect of TFs between the enhancer pair, divided by the union of TFs) per enhancer-enhancer correlation coefficient (N = 18,688). Fit line corresponds to a linear regression model with 95% confidence intervals.

**Supplementary Fig. 9.**
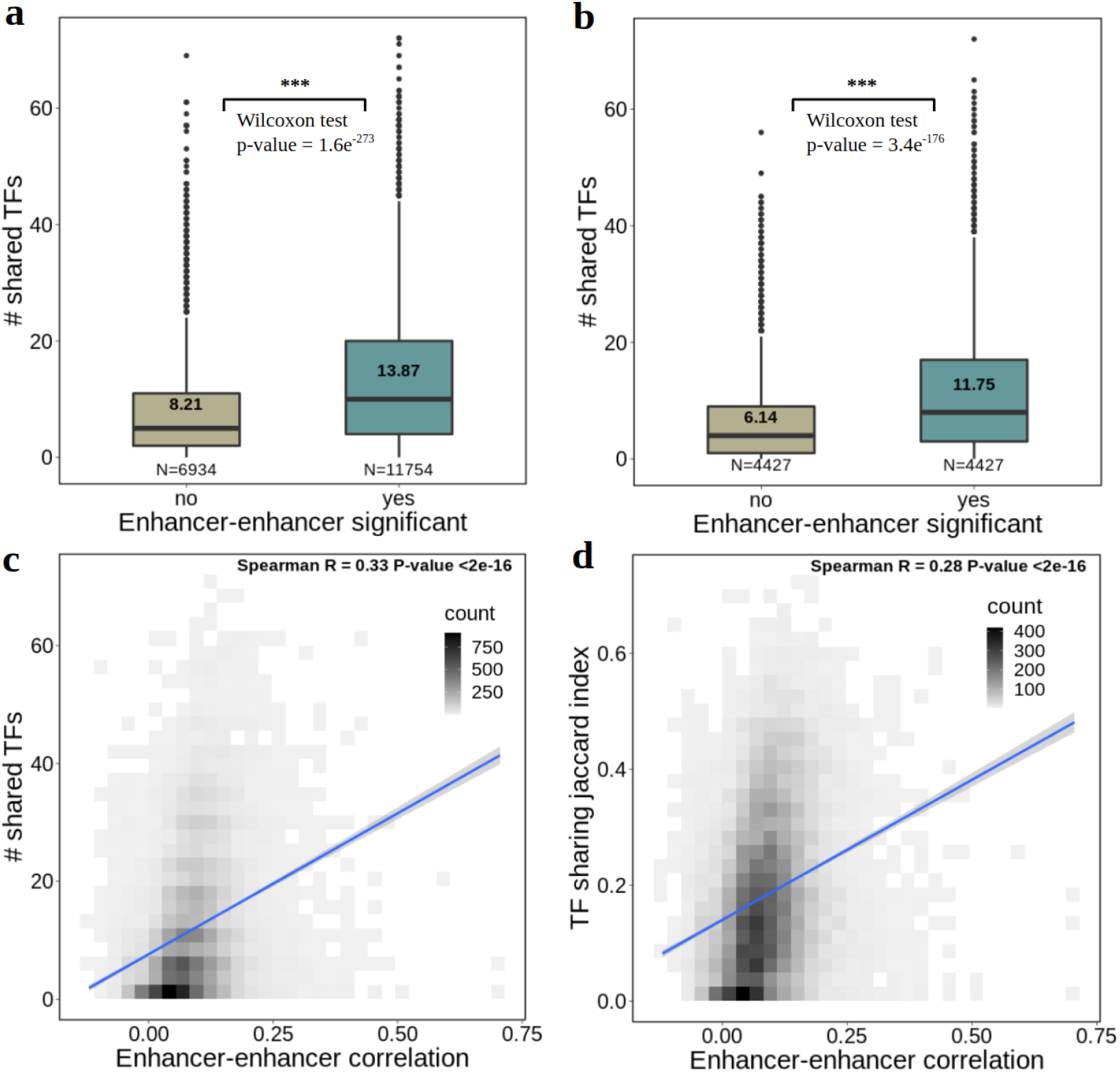
Transcription factor sharing on ABC enhancers using ReMap data. **a** Number of distinct TFs with binding sites in both enhancers of an enhancer-enhancer pair (shared TFs), depending on their association significance. The length of the box corresponds to the IQR with the centre line corresponding to the median, the upper and lower whiskers represent the largest or lowest value no further than 1.5 × IQR from the third and first quartile, respectively. Values above the median line represent the mean; **b** Same as previous, but significant and non-significant enhancer pairs are matched for distance (see Methods); **c** Number of shared TFs per enhancer-enhancer correlation coefficient (N = 18,688). Fit line corresponds to a linear regression model with 95% confidence intervals; **d** TF sharing Jaccard Index (intersect of TFs between the enhancer pair, divided by the union of TFs) per enhancer-enhancer correlation coefficient (N = 18,688). Fit line corresponds to a linear regression model with 95% confidence intervals.

**Supplementary Fig. 10.**
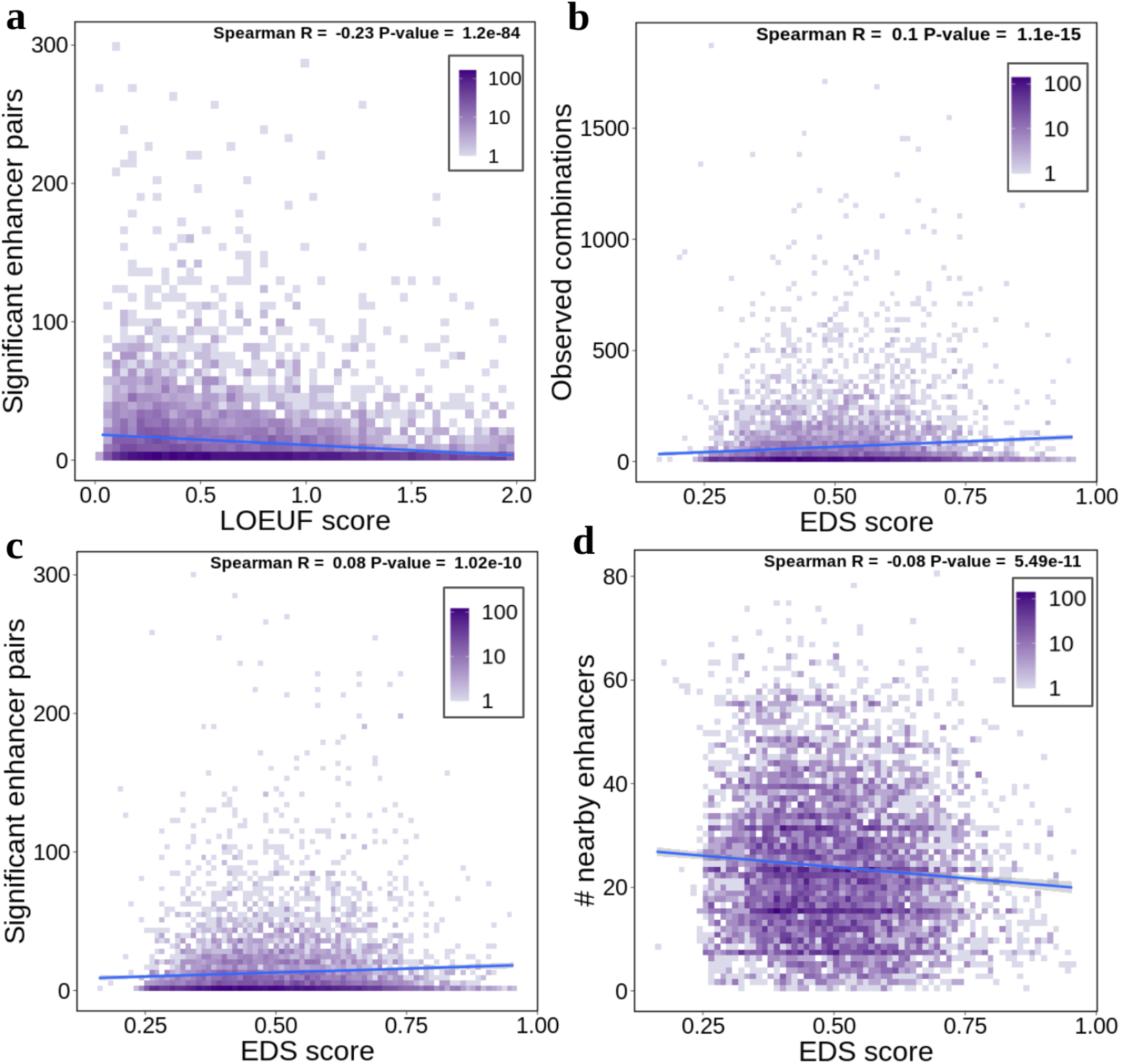
Enhancer correlation comparison with gene essentiality and enhancer redundancy scores. **a** number of significant enhancer-enhancer pairs and gene LOEUF score (N = 6895); **b** number of enhancer combinations observed in at least one cell per enhancer-domain score (EDS, N = 6925); **c** number of significant enhancer-enhancer pairs and EDS score; **d** number of enhancers within 1Mb of the gene TSS (regardless of gene-enhancer association significance) per EDS score. Fit lines correspond to a linear regression model.

